# Antibiotic resistance genes detected in lichens: insights from *Cladonia stellaris*

**DOI:** 10.1101/2025.05.28.656581

**Authors:** Marta Alonso-García, Paul B. L George, Samantha Leclerc, Marc Veillette, Caroline Duchaine, Juan Carlos Villarreal

## Abstract

**Background and Aims:** Antibiotics are natural compounds produced by microorganisms that have long existed in ecosystems. However, the widespread clinical and agricultural use of antibiotics has intensified selective pressures on bacteria, leading to the proliferation of antibiotic resistance genes (ARGs). The increasing prevalence of these genetic elements now poses a major global health threat. While ARGs are well documented in anthropogenically influenced environments, their distribution and origins in remote ecosystems, such as the boreal forests, remain poorly understood. Here, we investigate the occurrence, diversity, and potential origins of ARGs in the boreal lichen *Cladonia stellaris*.

**Methods:** We conducted the first targeted assessment of ARGs in lichens by analyzing 42 *C. stellaris* samples from northern and southern lichen woodlands (LWs) in eastern Canada. Using high-throughput quantitative PCR, we screened for 33 ARGs and three mobile genetic elements (MGEs), quantifying their relative abundance. Bacterial community composition was characterized via 16S rRNA gene sequencing. Statistical analyses evaluated geographical patterns, ARGs-taxa association, and the influence of latitude on ARG distribution.

**Key Results:** Ten ARGs conferring resistance to four antibiotic classes (aminoglycosides, beta-lactams, quinolones and sulfonamides), along with one MGE, were detected. Three ARGs, *blaCTX-M-1*, *qnrB*, and *qepA*, were highly prevalent, with *qepA* often surpassing 16S rRNA gene abundance. Latitude significantly influenced ARG profiles, whereas bacterial community composition did not. Network analysis identified *Connexibacter*, *Granulicella*, and *Novosphingobium* as potential hosts for *qnrB*, and *Tundrisphaera* and *Terriglobus* for *qepA*. To explain ARGs presence, we explored two hypotheses: bioaerosol dispersal from anthropogenic sources, and endogenous development through co-evolution between lichen-produced antimicrobial compounds and their associated bacterial communities.

**Conclusions:** Our findings demonstrate that *C. stellaris* harbors diverse ARGs in remote boreal ecosystems, highlighting the ecological complexity of ARG persistence and the need to investigate not only ARG presence, but also the processes driving their distribution in natural environments.

## Introduction

Antibiotics are a class of secondary metabolites naturally produced by microorganisms, such as bacteria or fungi, or chemically synthetized analogous compounds (Demain and Sanchez 2009). In nature, they serve various ecological roles, acting as signaling molecules and defense mechanisms. As signaling molecules, antibiotics are crucial for maintaining microbial balance and facilitating interactions within microbial communities (Henke and Bassler 2004; Linares *et al*. 2006; Schertzer *et al*. 2009). As defense mechanisms, they provide producing organisms with a competitive advantage by inhibiting the growth of rival microbes, thus securing resources and space (Waksman and Woodruff 1940). According to the arms-shield hypothesis, the production of antibiotics and the development of resistance to them is an ongoing evolutionary battle (Han *et al*. 2022). Some microorganisms have evolved the ability to produce antibiotics as a means of defense, while their competitors have developed resistance mechanisms through natural selection, enabling them to survive antibiotic exposure. This resistance capability is conferred by antibiotic resistance genes (ARGs), which are segments of DNA encoding proteins that enable bacteria to survive exposure to antibiotics. ARGs can develop intrinsically within the bacterial genome or acquired through horizontal gene transfer (HGT) (Thomas and Nielsen 2005; Barlow 2009) from other bacteria via mobile genetic elements (MGEs) such as plasmids (Partridge *et al*. 2018; Razavi *et al*. 2020; Ni *et al*. 2023), or via bacteriophages (Balcazar 2014; Penadés *et al*. 2015). Their existence in the environments predates the anthropogenic use of antibiotics (Wright 2007; Martínez 2008) with evolutionary evidence tracing their development from thousands (D’costa *et al*. 2011) to billions of years (Hall and Barlow 2004).

Since the 1940s, when antibiotics began to be used in medicine, their application has expanded to sectors such as agriculture (e.g. Anderson and Gottlieb, (1952); McManus et al., (2002)), aquaculture (e.g. Cabello, (2006); Chen et al., (2020)), and animal husbandry (e.g. Hong et al., (2013); Busch et al., (2020); Karwowska, (2024)). Such extensive, and at times inappropriate, use of antibiotics has led to the proliferation of ARGs and ARG-carrying bacteria in both natural and anthropogenic ecosystems. The World Health Organization has identified antibiotic resistance as one of the top public health threats of the 21^st^ century. In 2019 alone, over 4.95 million deaths were linked to antibiotic resistance, with approximately 1.27 million directly attributed to infections caused by antibiotic-resistant bacteria (Murray *et al*. 2022). At the national level, antimicrobial resistance was responsible for the deaths of approximately 5,400 Canadians in 2018. If current trends continue, resistance rates are projected to increase from 26% in 2018 to 40% and potentially up to 396,000 cumulative deaths by 2050, with economic losses ranging from 13 to 21 billion CAD (Council of Canadian Academies 2019).

The spread of ARGs in the environment occurs not only through local contamination near anthropogenic sources, such as the direct release of ARG-carrying bacteria via wastewater discharge or agricultural runoff (Almakki *et al*. 2019; Junaid *et al*. 2022), but also by atmospheric processes that enable their dispersal over much greater distances. Bioaerosols, which are airborne particles of biological origin encompassing bacteria, viruses, and fungi, can carry ARGs over distances ranging from meters to hundreds of kilometers (Brunet *et al*. 2017; Griffin *et al*. 2017). Bioaerosols are transported by atmospheric currents (Jin *et al*. 2022; Wang *et al*. 2022), before settling out of the air through dry deposition (gravity-driven settling onto surfaces) or precipitation into terrestrial or aquatic environments (Di Cesare *et al*. 2017; O’Malley *et al*. 2023). For example, clouds are increasingly recognized as dynamic microbial ecosystems, rich in viable microorganisms and genetic material, including ARGs (Rossi *et al*. 2022). Wildfires, while less studied as a vector of ARG dispersion, represent an additional mechanism by which large quantities of soil and plant-associated microorganisms can be aerosolized and transported through smoke (Moore *et al*. 2021). Despite the intense heat associated with wildfires, studies have shown that up to 80% of DNA-containing cells in smoke remain viable and metabolically active (Moore *et al*. 2021).

Lichen woodlands (LWs) provide a particularly valuable system for investigating the presence and distribution of ARGs. Characterized by open forests with minimal tree canopy, LWs offer little physical obstruction to airborne particles, allowing microorganisms suspended in the atmosphere to settle directly onto the ground layer. This ground layer is dominated by a continuous and exposed cover of lichens (Payette *et al*. 2001; Girard *et al*. 2017), which renowned for their ability to absorb and accumulate substances from their surrounding environment, including airborne particles. Interestingly, LWs also provide an opportunity to explore endogenous mechanisms of antibiotic resistance development. Lichens are known to produce a variety of antimicrobial compounds (Burkholder *et al*. 1944; Yilmaz *et al*. 2004; Rankovič *et al*. 2010; Shrestha and St. Clair 2013), which exert selective pressure on their associated bacterial communities (Grube *et al*. 2009; Grimm *et al*. 2021), and may promote the development of resistance, as proposed for the lichen species *Rhizocarpon geographicum* (Miral *et al*. 2022).

Across Canada, LWs cover approximately 2 million km^2^ of which, nearly 300,000 km^2^ can be found in the province of Quebec (Payette and Delwaide 2018), with *Cladonia stellaris* (Opiz) Pouzar & Vězda being the dominant and most representative lichen species. LWs are situated north of the closed-crown forest zone and south of the forest-tundra zone, between 52.00°N and 55.00°N, in remote regions characterized by low human population density and limited anthropogenic activities (Bhiry *et al*. 2011). An exception to the typical distribution of LWs occurs in Parc National des Grands-Jardins, Quebec (PNGJ; 47.68°N, 70.85°W), where a LW is found 500 km south of its usual range (Jasinski and Payette 2005). In 2023, the southern boundary between LWs and the closed-crown forest was severely affected by extreme wildfires (Gaboriau *et al*. 2023; Boulanger *et al*. 2024), during the most severe wildfire season on record, with over 4.5 million hectares of forest burned across Quebec (SOPFEU 2023). Although wildfires have long been a natural disturbance in boreal regions (Serge Payette 1992), the exceptional intensity of the 2023 fire season (Jain *et al*. 2024) raises new questions about their potential role in atmospheric dissemination of ARGs (Moore *et al*. 2021).

Considering the potential contribution of multiple atmospheric processes, including wildfire smoke, prevailing winds, precipitation and cloud dynamics, to the deposition of ARGs in LWs, and the lichens’ ability to accumulate airborne substances, we propose that lichens may act as reservoirs for ARGs. This capacity could also be reinforced by possible coevolutionary dynamics between lichen-associated bacteria and their host under selective pressures. To investigate this, we conducted the first targeted assessment of ARGs in lichen samples using high-throughput quantitative PCR (HT-qPCR). The objectives of this study are to i) test the presence of ARGs in lichens from both northern and southern LWs, ii) quantify their abundance, iii) compare the diversity and abundance of ARGs between northern and southern LWs, and iv) explore the relationship between their ARG profiles and constituent bacterial communities. To frame our investigation, we consider two main hypotheses to explain the origins of bacteria-carrying ARGs in *C. stellaris*: first, that long-distance dispersal by bioaerosols introduces ARGs to LWs (Kormos *et al*. 2022; George *et al*. 2022), by transporting bacteria-carrying ARGs from urban and agricultural environments where antibiotics are frequently used; and second, that ARGs in lichen-associated communities may be indigenous, developing as a response to selective pressures imposed by the antimicrobial compounds produced by lichens (Francolini *et al*. 2004; Goga *et al*. 2021; Zorrilla *et al*. 2022). This latter scenario, consistent with the arms-shields race hypothesis, suggests that the evolutionary interplay between the lichen host and its associated bacteria may have driven the development of ARGs as an adaptive strategy.

## Material and methods

### Study areas and sampling

This study builds upon previous data collected by Alonso-García et al. (2021) and Alonso-García and Villarreal A. (2022), which investigated the factors shaping the bacterial community composition of lichens across Quebec, Canada. *Cladonia stellaris* samples were collected from two location: the first near Kuujjuarapik-Whapmagoostui (55.28,-77.75) in the north and the second from PNGJ (47.68,-70.85) in the south. Kuujjuarapik-Whapmagoostui has a mean annual temperature of-3.6°C and a mean annual precipitation of 640 mm (Prairie Climate Centre University of Winnipeg 2022). In this region, LWs develop on sandy terraces and wind-affected areas, where *Picea glauca* (Moench) Voss and *Picea mariana* (Mill.) Britton, Sterns & Poggenb. coexist with a ground cover dominated by *C. stellaris* and ericaceous shrubs such *Betula glandulosa* Michx., *Rhododendron groenlandicum* Retzius, and *Salix glauca* L. (Bhiry *et al*. 2011). These open forests grow on acidic, nutrient-poor, well-drained soils (Payette 1993; Payette *et al*. 2001).

The PNGJ has a mean annual temperature of 2°C, and an annual precipitation of approximately 1200 mm (Payette *et al*. 2000; Prairie Climate Centre University of Winnipeg 2022). The park features a mosaic of closed-crown spruce-moss forests and open-crown lichen woodlands dominated by *P. mariana*, Ericaceae species and *C. stellaris* (Jasinski and Payette 2005), thriving on acidic, nutrient-depleted moraine-derived soils and granitic outcrops (S. Payette 1992; Payette and Morneau 1993). Although both regions are relatively remote compared to urban centers, potential sources of anthropogenic contamination exist near the study sites. In Kuujjuarapik-Whapmagoostui, there is a possibility that domestic waste, healthcare services, and potential wastewater seepage may introduce antibiotics into nearby ecosystems. The PNGJ, has protected status, restricting human activities. However, it is located within the Charlevoix region, where agriculture and livestock farming are widespread (MRC de Charlevoix 2022), and its proximity to the town of Baie-Saint-Paul (less than 30 km away), could expose the park to anthropogenic contaminants.

Sampling was performed in 2018 using sterilized steel forceps, targeting thalli of *C. stellaris*, which were immediately stored at-20°C. Further methodological details are available in Alonso-García et al., (2021); and Alonso-García and Villarreal A., (2022). In the present study, 42 samples were selected for analyses: 18 from seven sites around Kuujjuarapik-Whapmagoostui (northern LW), and 24 from PNGJ (southern LW) (Table S1).

### Data processing and analysis of 16S rRNA genes

To profile bacterial community composition, we used amplicons of V3-V4 region of the 16S rRNA gene generated by Alonso-García and Villarreal A., (2022). These data were passed through an established DADA2 pipeline (Callahan *et al*. 2016) in the R environment (R Core Team 2021) for processing amplicon sequence variants (ASVs, Callahan et al. (2017)). We transformed ASV counts to relative abundances using the phyloseq package (McMurdie and Holmes 2013) for comparisons of biodiversity between northern and southern LWs. To calculate the Shannon diversity index, we first normalized the ASV table using the DESeq2 package (Love *et al*. 2014), applying the poscounts method. This approach estimates size factors using the median-of-ratios method while excluding zero counts, making it particularly suitable for sparse amplicon data typical of microbial community profiles. We tested for differences in bacterial alpha diversity between LWs using an unpaired t-tests, after confirming normality and homogeneity of variances using the Shapiro-Wilk and Levene’s tests, respectively. To analyze beta diversity, we built a Bray-Curtis distance matrix based on relative abundances and performed a PCoA (Principal Coordinates Analyses). We assessed differences in bacterial community composition between LWs with PERMANOVA (Permutational Multivariate Analysis of Variance) test, preceded by a check of the homogeneity of dispersions using the betadisper function from the vegan package (Oksanen *et al*. 2024). We used the Bray-Curtis distance matrix from the relative abundance data to perform db-RDA to evaluate the influence of the latitude of LW on the bacterial community composition, fitting the model with the capscale function in vegan package (Oksanen *et al*. 2024), with LW as explanatory variable. The significance of the db-RDA model was assessed using permutation tests.

### SmartChip high-throughput quantitative PCR

Bacterial biomass was assessed via qPCR using the 16S rRNA marker gene (primers and probe as described in Bach et al. (2002). The qPCR was conducted on a Bio-Rad CFX-384 Touch™ Real-Time PCR Detection System (Bio-Rad, Montreal, CA) under the following thermoprotocol: 95 °C for 3 min; followed by 40 cycles of 95 °C for 20 s and 62 °C for 1 min. Results were validated only if the accompanying standard curves showed efficiency values between 90% and 110%.

We employed the SmartChip Real-Time PCR System (Takara Bio USA, Inc.) to screen for the presence of ARGs and MRGs. The HT-qPCR was performed following the manufacturer’s protocol. Each 250 μl reaction contained the appropriate primers for the targeted ARGs, along with the lichen DNA extracted previously in Alonso-García and Villarreal A. (2022).The selection of ARGs was guided by the framework established George et al. (2022), ensuring comparability with previous global studies. We targeted 33 ARGs and three MGEs (Table 1). We included positive controls consisting of synthetic target gene sequences at concentrations of 10[, 10³, and 10¹ copies per microliter, along with negative controls (no-template controls), to verify specificity and the absence of contamination. Each sample was analyzed in triplicate. A gene was considered present if detected in at least two out of the three technical replicates. The Ct values from these replicates were averaged for further analysis.

**Table 1.**
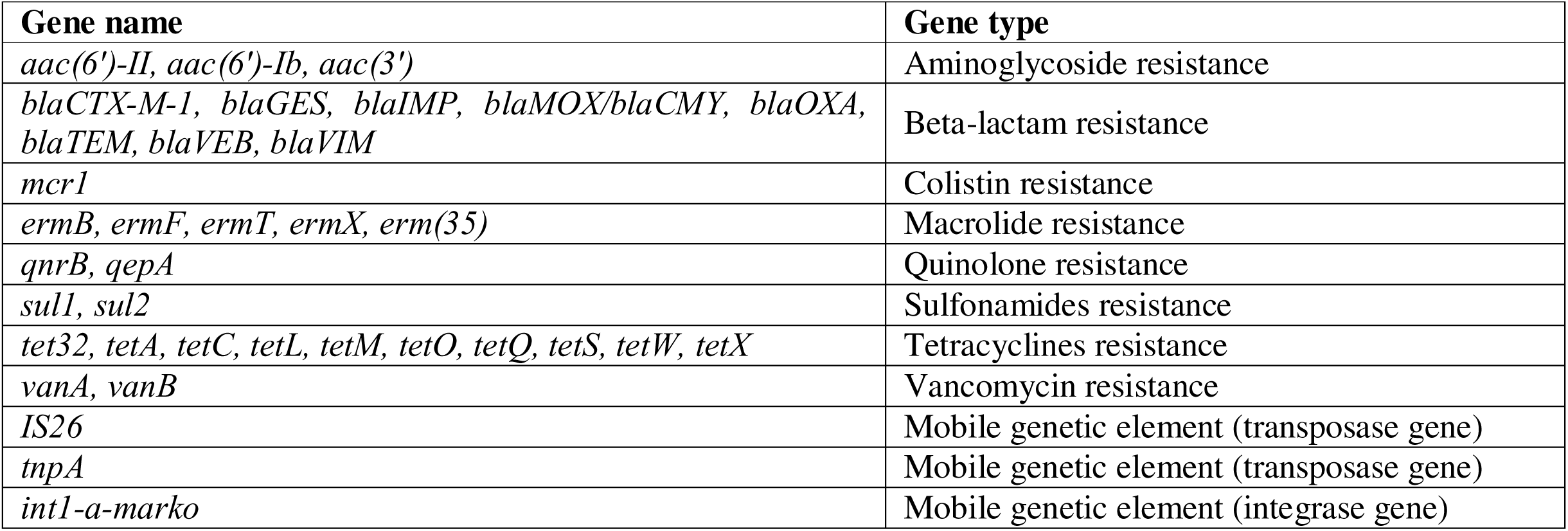
List of gene targets used for HT-qPCR analyses.

The relative abundance of each ARG/MGE compared with 16S rRNA was calculated using the 2^-ΔCt^ method, where ΔCt = Ct(ARG or MGE) - Ct(16S rRNA); hereafter referred to as ARG relative abundance. To visualize the variation of ARGs across the lichen samples, we created a heat map using the relative abundance values. A logarithmic transformation (log) was applied to the abundance values to better represent the wide range of abundances in the heat map. We compared ARG relative abundance between northern and southern LWs using exclusively ARGs with a prevalence greater than 50% across all samples. To test for statistical differences in ARG abundance between northern and southern LWs, we performed Wilcoxon rank-sum tests with Benjamini-Hochberg (BH) correction due to non-normal distribution of the data revealed by Shapiro-Wilk test. Prior to testing, abundance values of zero were excluded from the dataset to avoid skewing the results. Additionally, we assessed the overall ARG abundance in each lichen sample, by summing the relative abundances of all ARGs for each sample, and we applied a Wilcoxon rank-sum test to compare it between the two LWs. We assessed the influence of the latitude of LW on the relative abundance of the eleven ARGs detected in *C. stellaris* performing a db-RDA on the Bray-Curtis distance matrix of ARG relative abundance as described above.

### Comparison of ARGs and bacterial communities in lichens

We assessed how much of the variance in the relative abundance of ARGs could be explained by the relative abundance of bacterial families. The relative abundances of ARGs served as the response variables, while the relative abundances of bacterial families were the explanatory variables. To check for multicollinearity among the explanatory variables, we calculated Spearman’s correlation coefficients. We used a threshold of 0.4 for correlations to identify and remove highly co-correlated bacterial families from the analysis. The removed families were: Bdellovibrionaceae, Sphingomonadaceae, Isosphaeraceae, SM2D12, Caulobacteraceae, and Microbacteriaceae. Given that the linearity assumption between ARGs and bacterial families was not confirmed, we opted to use Canonical Correspondence Analysis (CCA), which can handle non-linear relationships. The significance of the CCA model was tested using ANOVA with permutation.

To explore potential relationships between ARGs and bacterial taxa associated, we conducted network analyses. First, we prepared a dataset combined ARG relative abundances with bacterial genera abundances, transformed using centered log-ratio to standardize the data. We calculated Spearman correlation coefficients (|ρ|) and corresponding p-values using the rcorr function from the Hmisc package in R (Harrell J. 2024). We set significance thresholds at a correlation coefficient greater than 0.2 and an adjusted p-value below 0.05. To control for multiple testing, we applied BH correction to the p-values. Significant correlations were retained, and their values were used to construct the network, where nodes represent ARGs or bacterial taxa, and edges represent significant correlations between them. The network was generated using the igraph package in R (Csardi and Nepusz 2006), with edge weights corresponding to the absolute values of the Spearman correlation coefficients. The direction of the correlation (positive or negative) was annotated, and correlation strength was categorized as weak, moderate, strong or very strong based on the absolute value of the correlation coefficient (ranges: 0.2-0.4, 0.4-0.6, 0.6-0.8, 0.8-1.0, respectively). For further visualization and editing, we exported the network data, including node attributes and edges with adjusted p-values, to a CSV file compatible with Cytoscape software (Shannon *et al*. 2003).

## Results

### Bacterial community differs between LW

A total of 481 unique ASVs were retained after quality and prevalence filters from 48 lichen samples. The ASVs were assembled into ten phyla. Proteobacteria was the most abundant, accounting for 242 ASVs. The next most abundant phyla were Acidobacteriota (86 ASVs), Planctomycetota (78 ASVs), followed by Verrucomicrobiota (42 ASVs) (Figure S1A). Among the ASVs assigned to genera, *Tundrisphaera* (65 ASVs), *Granulicella* (41 ASVs), and LD29 (33 ASVs) were the most abundant, while genera such as *Conexibacter*, *Terriglobus*, and *Novosphingobium* were represented by only one or two ASVs each (Figure S1B, Table S2). Unpaired t-test revealed that alpha diversity was higher in northern than in southern LWs (p = 0.0001) (Figure 1A). The Bray-Curtis distance based PCoA showed that bacterial communities of *C. stellaris* differ between LWs, with no overlap, with axes 1 and 2 explaining 22% and 10.5% of the total variance, respectively (Figure **1**. **A** 1B). PERMANOVA results confirmed that the separation between LWs was statistically significant (R = 0.143, p = 0.001). The db-RDA model revealed that the latitude of LW explained 14.23% of the variance in the bacterial community composition. The global permutation test confirmed that the model was statistically significant (F = 6.634, p = 0.001).

**Figure 1.**
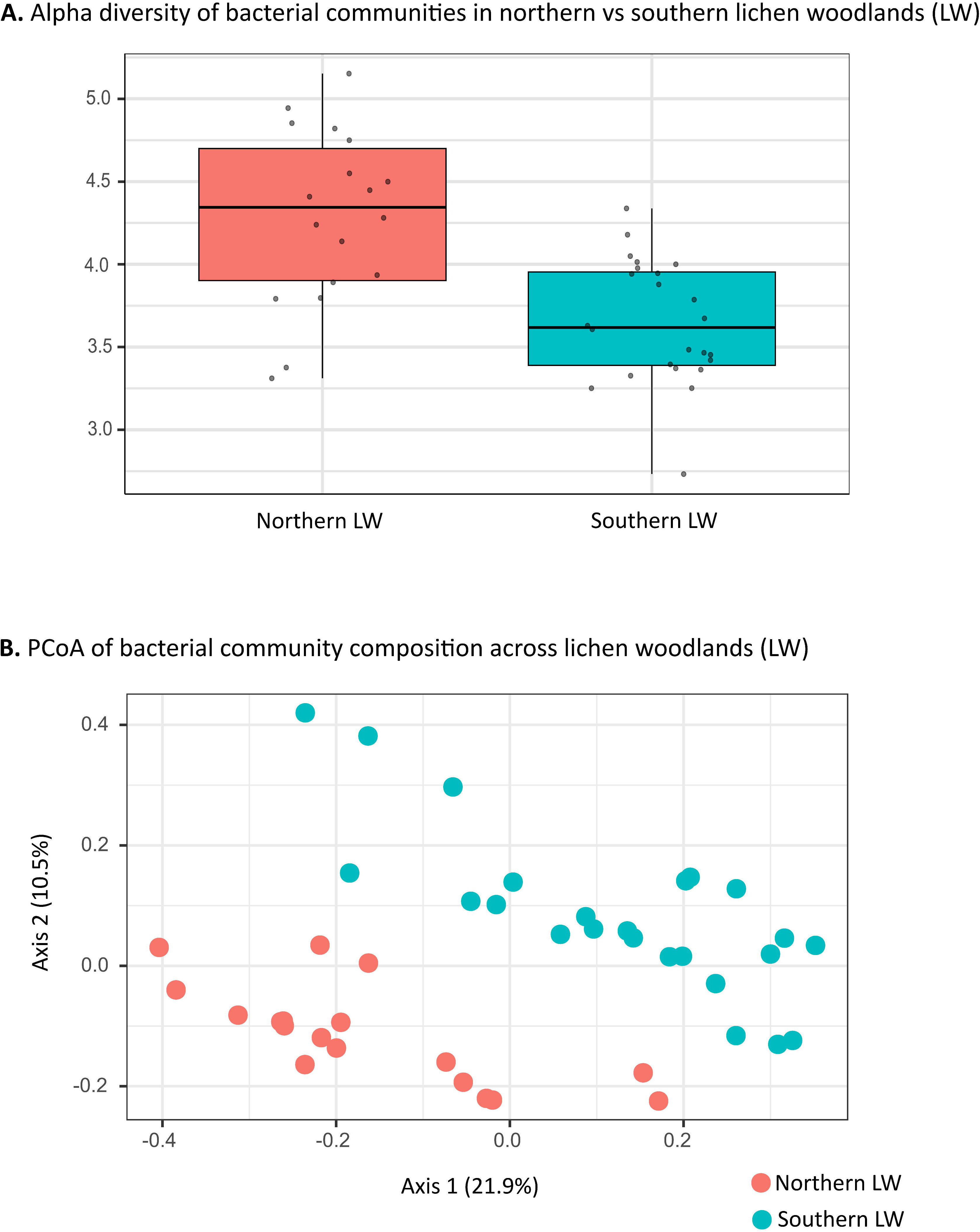
A. Alpha diversity Shannon index of bacteria associated with *Cladonia stellaris* from northern and southern lichen woodlands (LW). The median (horizontal line), quartiles (edges of the box) and 1.5X the interquartile range of (whiskers) are displayed. Points represent individual observations. Student’s t-test results indicated a significant difference in alpha diversity between the two LWs (p-value < 0.05). **B.** Principal coordinates analysis (PCoA) of bacteria associated with *Cladonia stellaris* from northern and southern LW. Samples are colored according to LW latitude. PERMANOVA results indicated a significant difference in the bacterial community composition between the two LWs (R = 0.14303, p-value < 0.01).

### Characterisation of ARGs present in lichens

Among the 33 ARGs and three MGEs tested, ten ARGs and one MGE were detected in *C. stellaris* samples, corresponding to resistance to aminoglycosides (*aac(6’)-Ib* and *aac(3’)*), beta-lactams (*blaCTX-M-1*, *blaMOX/CMY*, *blaTEM* and *blaVIM*), quinolones (*qepA*, and *qnrB*), and sulfonamides (*sul1* and *sul2*), as well as one MGE (*int1-a-marko*). The relative abundances of the detected ARGs and MGE are shown in Figure 2 and Table S3. While most ARGs and the MGEs exhibited lower abundance compared to the 16S rRNA gene, *qepA* was an exception, displaying higher relative abundance in most samples (Figure 2). Three ARGs, *blaCTX-M-1*, *qnrB* and *qepA*, were the most prevalent, each occurring in more than 50% of the samples. The *blaCTX-M-1* gene, conferring beta-lactamase resistance, was found in 35 samples (16 from the northern LW and 19 from the southern LW). The quinolone resistance genes *qnrB* and *qepA* were detected in 34 (14 northern LW, 20 southern LW) and 23 (14 northern LW, 9 southern LW) samples, respectively (Figure 2, Table S3). Comparisons of relative abundance of the three prevalent ARGs between northern and southern LWs (Figure 3) revealed a significant difference for *qnrB* (adjusted p = 0.027), with higher abundance in southern LW. However, when considering the total relative abundance of all detected ARGs combined (Figure S2), no significant differences were found between the two regions (adjusted p = 0.37). Finally, the db-RDA analysis revealed that the latitude of LW explained 12.61% of the variance in the relative abundance of ARGs associated with *C. stellaris*. The global permutation test confirmed that the model was statistically significant (F = 5.696, p-value = 0.005).

**Figure 2.**
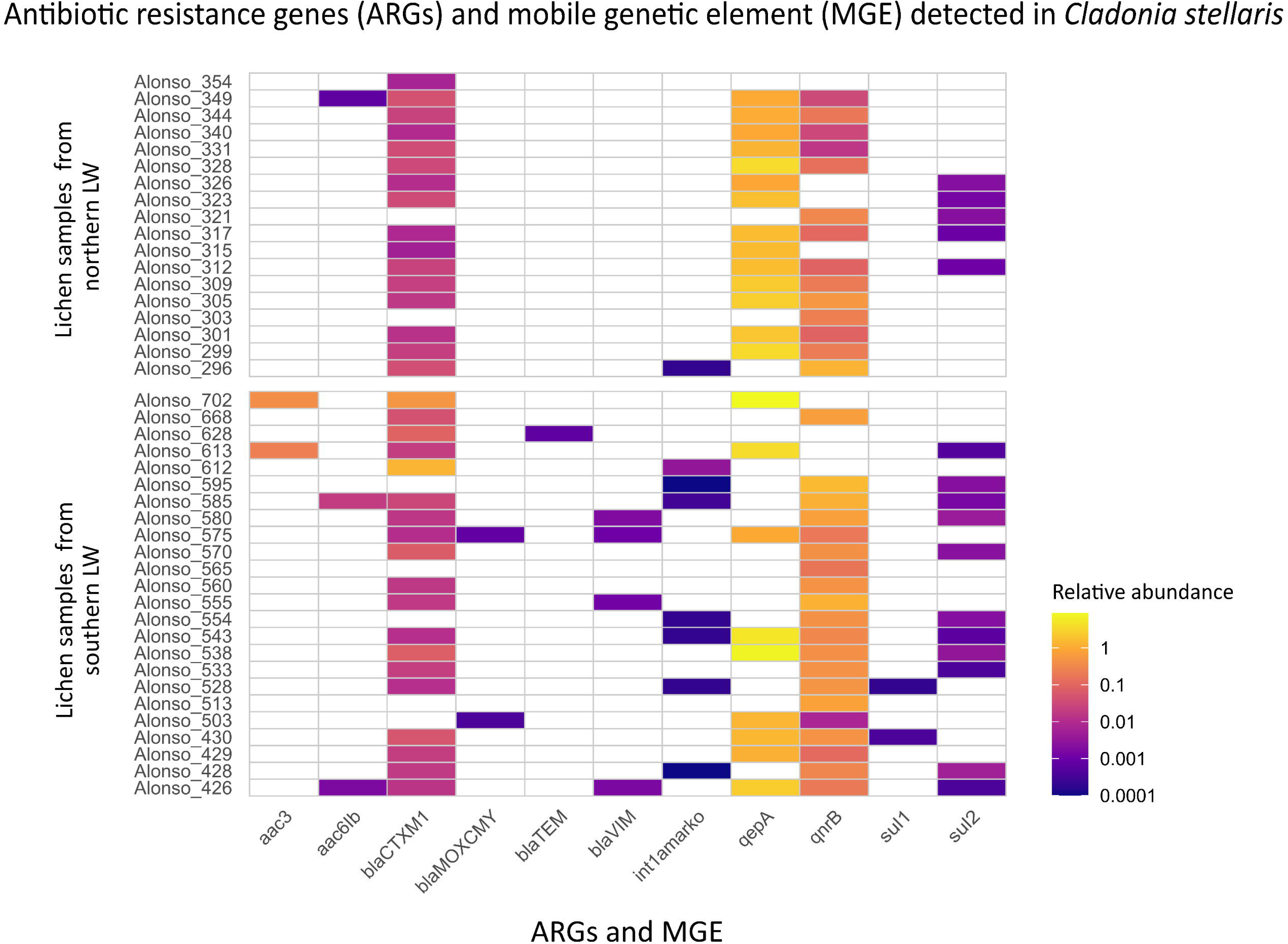
Heat Map of ten antibiotic resistance genes (ARGs) and one mobile genetic element (MGE) found in *Cladonia stellaris* samples. Each tile represents the logarithmic (log) abundance of a specific ARG/MGE in each lichen sample, with darker colors indicating higher abundance and white tiles indicating an absence of the gene. Samples are grouped by lichen woodland (LW) latitude, on the y-axis, while the ARGs/MGE are displayed on the x-axis.

**Figure 3.**
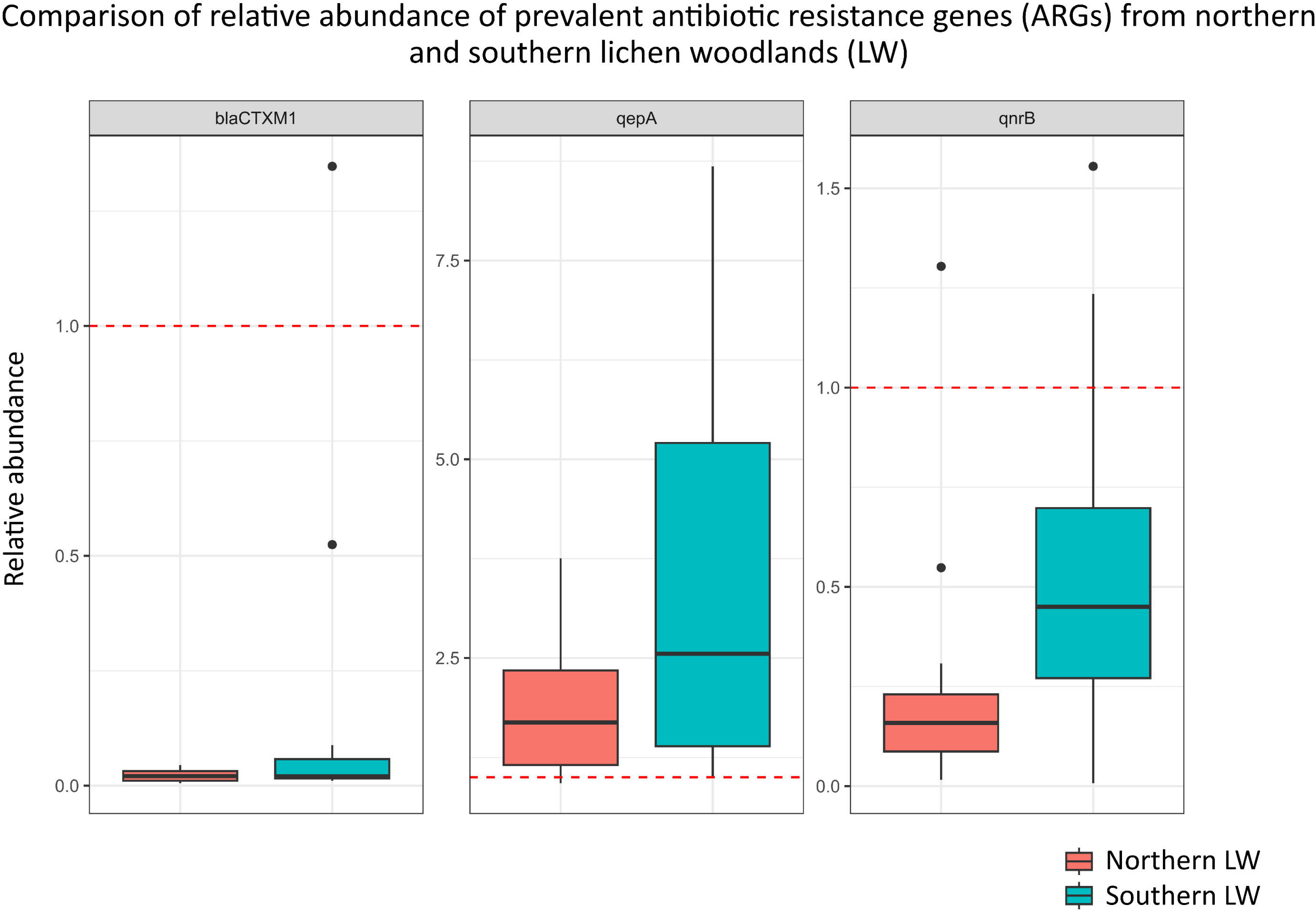
Relative abundance of three prevalent antibiotic resistance genes (ARGs) in *Cladonia stellaris* from northern and southern lichen woodlands (LWs). The median (horizontal line), quartiles (edges of the box) and 1.5X the interquartile range of (whiskers) are displayed. Points represent individual observations. A dashed red horizontal line at y = 1 indicates the reference value, where values equal to 1 represent abundance equivalent to the 16S rRNA gene. Values greater than 1 indicate higher abundance compared to the 16S rRNA gene, while values less than 1 indicate lower abundance. Wilcoxon rank-sum results indicated significant differences in the relative abundance of *qnrB* between the northern and southern LWs (adjusted p-value = 0.0274).

### Relationship between ARGs and bacterial communities in lichens

The CCA did not yield statistically significant results (F = 1.7, p = 0.139), indicating that the relative abundances of bacterial families did not account for a significant proportion of the variance observed in ARG relative abundances. Network analysis displayed the relationships between bacterial genera and the two quinolone ARGs (*qnrB* and *qepA*), with no connexions observed for *blaCTX-M-1*. The overall network comprised 32 nodes and 147 edges, with an average of 9.19 neighbors per node, a clustering coefficient of 0.527, and a network density of 0.296 (Table 2). These metrics indicate a moderately interconnected network with a cohesive structure, where no bacterial genera or ARGs were isolated (Figure 4).

**Figure 4.**
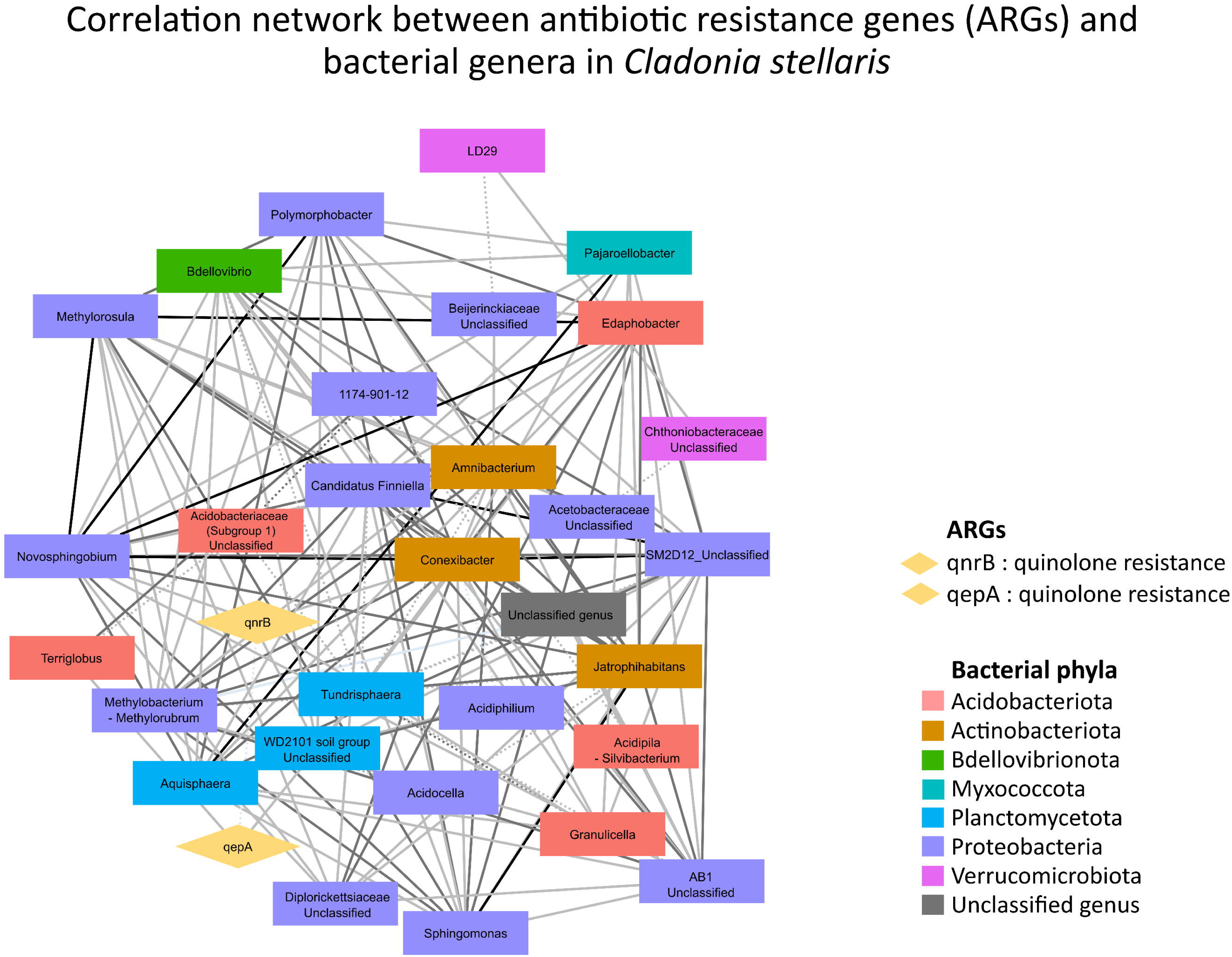
Network analysis of correlations between antibiotic resistance genes (ARGs) and bacterial genera associated with *Cladonia stellaris*. The network was constructed using Spearman correlation coefficients, displaying significant correlations (|ρ| > 0.2 and adjusted p-value < 0.05) between ARGs and bacterial genera. Nodes represent ARGs (diamonds) or bacterial genera (rectangles), with edges depict significant correlations. Edge color indicates correlation strength, categorized as weak (0.2 < |ρ| ≤ 0.4), moderate (0.4 < |ρ| ≤ 0.6), or strong (|ρ| > 0.6). Solid edges indicate positive correlation, and dotted edges represent negative correlations.

**Table 2.**
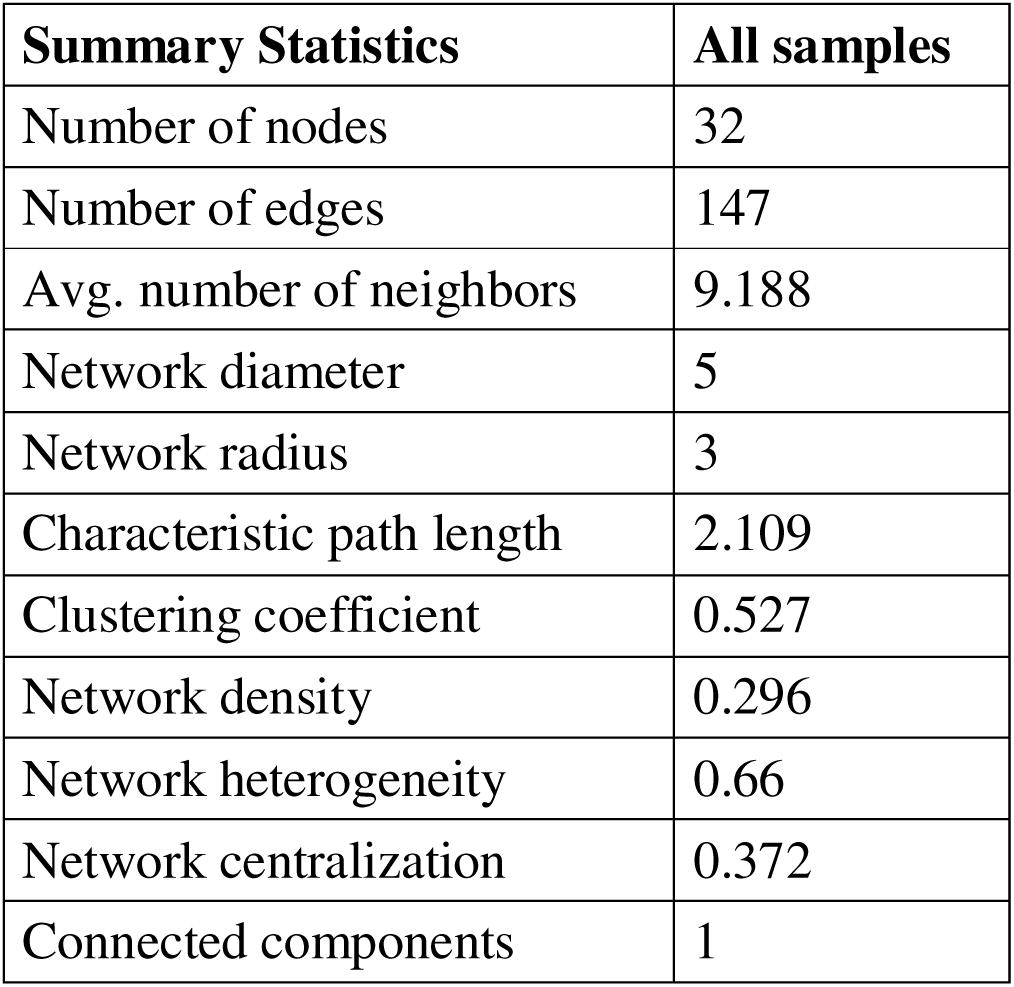
Summary of key network metrics of bacterial communities and antibiotic resistance genes (ARGs) associated with *Cladonia stellaris*.

Among the genera identified, *Conexibacter* (represented by 2 ASVs) emerged as a key node within the bacterial community of *C. stellaris*. It exhibited the highest number of connections (20) and high centrality indices (closeness centrality = 0.67, betweenness centrality = 0.219) (Figure 4, Table S4). This genus was positively correlated with *qnrB* (|ρ| = 0.54). *Granulicella* (41 ASVs) and *Novosphingobium* (1 ASV), though differing in abundance, showed similar patterns in the network. Both genera showed moderate positive correlations with *qnrB* (|ρ| = 0.54 for *Granulicella* and |ρ| = 0.43 for *Novosphingobium*) and had relatively low betweenness centrality (0.010 for *Granulicella* and 0.027 for *Novosphingobium*). Additionally, *Granulicella* contributed eight connections to the network, indicating a moderate role in the network structure, while *Novosphingobium* had 16 connections, reflecting a stronger presence in the network (Figure 4, Table S4). Two different genera were positively correlated with *qepA*: *Tundrisphaera* (|ρ| = 0.43) and *Terriglobus* (|ρ| = 0.45). While *Tundrisphaera* (65 ASVs) had 11 connections, *Terriglobus* (1 ASV) had only three. Both genera displayed relatively low betweenness centrality (0.089 for *Tundrisphaera* and 0.007 for *Terriglobus*) (Figure 4, Table S4), suggesting a more localized role in linking different parts of the network. A weak but significant negative correlation was observed between *qnrB* and *qepA* (|ρ| =-0.40) (Figure 4, Table S4), which may indicate that these two resistance genes do not co-occur frequently in the same bacterial community. However, given that *qepA* exhibits much higher relative abundance across samples compared to *qnrB*, this negative correlation may also reflect the influence of their relative abundance differences, which could affect their co-occurrence patterns.

## Discussion

Bacteria associated with *C. stellaris* from southern and northern LWs in Quebec harbored ten genes conferring resistance to four classes of antibiotics: aminoglycosides (*aac(6’)-Ib* and *aac(3’)*), beta-lactams (*blaCTX-M-1*, *blaMOX/CMY*, *blaTEM* and *blaVIM*), quinolones (*qepA*, and *qnrB*), and sulfonamides (*sul1* and *sul2*), as well as one MGE (*int1-a-marko*). One beta-lactam resistance gene (*blaCTX-M-1*) and two quinolones (*qepA*, and *qnrB*) were detected in over 50% of the samples, with *qepA* showing a particularly high relative abundance, surpassing the abundance of 16S rRNA gene in most samples. Comparisons between northern and southern LWs indicated a trend toward higher ARG relative abundance in southern samples; however, only the *qnrB* gene showed a statistically significant difference. Redundancy analyses revealed that LW latitude explained almost 13% of the variance in ARG profiles, suggesting a spatial effect. In contrast, no significant influence of bacterial family composition on ARG abundance was detected. Additionally, we identified the bacterial genera *Granulicella*, *Connexibacter*, and *Novosphingobium* as potential hosts for *qnrB*, and *Tundrisphaera* and *Terriglobus* for *qepA*. Although our data confirmed the presence of ARGs in *C. stellaris* across both northern and southern LWs, the mechanisms underlying the arrival and persistence of ARG in these ecosystems remain unclear. In the following sections, we explore two main hypotheses: i) long-distance dispersal via bioaerosols, and ii) endogenous development driven by coevolutionary dynamics between the lichen host and its associated bacteria (arms-shields race hypothesis). These interpretations remain preliminary, and further research will be essential to address the unresolved questions raised by our findings.

### Influence of LW latitude on bacterial community and ARGs distribution

Consistent with the findings of Alonso-García and Villarreal A., (2022), our results revealed significant differences in the diversity and composition of bacterial communities associated with *C. stellaris* between northern and southern LWs, with northern LWs exhibiting higher bacterial diversity. Latitude accounted for around 14% of the variation in the bacterial community of lichen samples and about 13% of the variation in ARG relative abundance, suggesting that local factors specific to each LW influence both bacterial community and ARG abundance. While the impact of the local abiotic factors on bacterial community composition is well established (Cardinale *et al*. 2012; Klarenberg *et al*. 2020; Paulsen *et al*. 2024), recent studies have also demonstrated its influence shaping ARGs abundance and distribution. Soil physicochemical properties have been shown to structure soil ARG profiles (Song *et al*. 2021; Wang *et al*. 2024). In addition to abiotic factors, biotic drivers such as microbial community composition can also influence ARGs abundance (Bahram *et al*. 2018; Yan *et al*. 2021), although the strength and direction of this effect vary by habitat type. For instance, bacterial community structure influences ARG abundance in soils, whereas no such effect was observed in the phyllosphere of plants (Xiang *et al*. 2020). Our results are consistent with this complexity: while LW latitude explained a small but significant portion of the variation in ARG abundance, bacterial community composition itself had no detectable influence on abundance of those genes, suggesting that additional, yet unidentified, factors are also driving ARG abundance in LWs.

### Frequent and abundant ARGs: beta-lactams and quinolones resistance

#### Beta-lactam resistance in lichen-associated bacteria

Beta-lactams antibiotics are among the most widely used antimicrobial agents and are naturally produced by certain microorganisms, such as the fungi *Penicillium chrysogenum* or the bacteria *Agrobacterium radiobacter* (Sykes *et al*. 1981). In this study, we detected four beta-lactamase resistance genes (*blaMOX/CMY*, *blaTEM*, *blaVIM* and *blaCTX-M-1*) in *C. stellaris* samples. Three of these genes were sporadically detected, exclusively in the southern LW. This sporadic detection may be related to nearby anthropogenic activities, such as agriculture and livestock farming, as beta-lactam ARGs are frequently enriched in agricultural and urban environments (Chen *et al*. 2018; Yan *et al*. 2019; Xiang *et al*. 2020). Bioaerosol transport from these areas could contribute to their subsequent deposition in PNGJ (Bai *et al*. 2022; George *et al*. 2022; Yang *et al*. 2023).

Conversely, *blaCTX-M-1* was by far the most prevalent ARG, found in 83% of the *C. stellaris* samples. We suggest that *blaCTX-M-1* may be stably integrated within the lichen-associated microbiome. Although lichens are not known to produce beta-lactams, they synthesize a wide array of antimicrobial compounds (Burkholder *et al*. 1944; Aslan *et al*. 2006; Rankovič *et al*. 2010; Mitrović *et al*. 2011; Shrestha and St. Clair 2013; Kosanić *et al*. 2014; Kosanić and Rankovic 2015), which could exert selective pressures favoring ARG retention over evolutionary timescales. Noël et al. (2021) demonstrated that lichen-associated bacteria developed mechanisms to survive in the presence of antibacterial compounds produced by lichens. Similarly, Agersø et al. (2019) suggested that some ARGs may be intrinsic to bacteria and originated from ancient resistomes rather than recent contamination. Beyond these ecological pressures, the widespread occurrence of *blaCTX-M-1* may also reflect its ability to persist independently of specific bacterial hosts. Indeed, no significant correlations were observed between this ARG and specific bacterial genera, a pattern also reported for other ARGs such as *blaTEM* (Jang *et al*. 2022) and *aac-(6)-Ib* (Yan *et al*. 2017). This lack of association could be explained by the frequent linkage of *blaCTX-M-1* to MGEs (Cantón *et al*. 2012; Partridge *et al*. 2018; Cormier *et al*. 2022), which promote HGT among diverse bacterial taxa. These mechanisms may allow *blaCTX-M-1* to disseminate broadly within microbial communities, independently of host identity, and to persist even as bacterial community composition shifts (Zhu *et al*. 2017; Li *et al*. 2023), as observed here between northern and southern LWs. Thus, the sporadic detection of beta-lactam *blaMOX/CMY*, *blaTEM* and *blaVIM* ARGs may reflect recent introductions through bioaerosols, while *blaCTX-M-1* presence might be explained by a combination of historical coevolutionary dynamics and the role of MGE, which facilitate its transfer across diverse bacterial hosts.

#### Quinolones resistance in lichen-associated bacteria

Quinolone are synthetic antibiotics introduced in the 1960s, with widespread use established by the 1980s (Hooper and Jacoby 2015). They are extensively employed in human and veterinary medicine, as well as in food production (World Health Organization 2018), contributing to growing contamination of natural environments (Cattoir *et al*. 2008; Ben *et al*. 2019; Zhai *et al*. 2024). In this study, we detected two quinolone resistance genes in *C. stellaris* samples: *qnrB* and *qepA*.

The *qnrB* gene exhibited a particularly high prevalence, being present in 81% of the lichen samples. It showed significantly higher relative abundance in southern lichens, pointing toward potential introductions via long-distance bioaerosol dispersal (Pilote *et al*. 2019; George *et al*. 2022) from anthropogenically influenced areas such as nearby agricultural and urban centers from the south. Our results further revealed that *qnrB* was positively associated with *Connexibacter*, a bacterial genus known for its resistance to quinolones (Monciardini *et al*. 2003), which exhibited the highest connectivity within the lichen bacterial network. *Connexibacter* may therefore act as an active reservoir of *qnrB*, facilitating its dissemination among other bacterial taxa. In addition, *Granulicella* and *Novosphingobium*, although less central within the network, were also identified as potential bacterial hosts for *qnrB*, consistent with previous reports linking *Novosphingobium* to other *qnr* genes (Yan *et al*. 2017). These findings suggest that multiple bacterial lineages may contribute to the persistence and circulation of *qnrB* within the *C. stellaris* microbiome. Furthermore, consistent with its potential mobility, *qnrB* frequently co-occurred with *blaCTX-M-1*, with both genes detected together in 77% of the *C. stellaris* samples, supporting previous reports of their potential genetic linkage on MGEs (Juraschek *et al*. 2022). While *blaCTX-M-1* may represent a historically embedded component of the lichen resistome (as proposed above), its association with MGEs could promote horizontal transfer. Such linkage could facilitate the simultaneous spread of resistance to beta-lactams and quinolones within bacterial communities associated with lichens. According to these findings, we suggest that the higher relative abundance of *qnrB* in southern lichens may reflect introductions through bioaerosol dispersal from anthropogenically influenced areas. Once introduced, the frequent co-occurrence of *qnrB* with *blaCTX-M-1* on MGEs (Juraschek *et al*. 2022) and its association with the central bacteria genus *Connexibacter* could facilitate its persistence within the lichen microbiome.

The *qepA* gene, encoding a quinolone efflux pump, was detected in 57% of the *C. stellaris* samples, with no significant differences in relative abundance between northern and southern LWs. Notably, *qepA* exhibited high relative abundance, often exceeding that of the 16S rRNA gene. We propose that the high abundance of *qepA* may reflect its ecological importance for the lichen-associated bacterial community. Efflux pumps serve diverse physiological roles for bacteria, particularly detoxification of potentially harmful compounds (Piddock 2006; Martínez *et al*. 2009; García-León *et al*. 2014). For instance, Alonso et al. (1999) demonstrated that environmental isolates of *Pseudomonas aeruginosa* collected prior to the widespread use of synthetic antibiotics could already actively expel quinolones, suggesting these mechanisms originally evolved against naturally occurring compounds. In the context of lichen symbioses, efflux pumps like *qepA* may provide a dual advantage: specific resistance against quinolones or quinolone-like compounds that might be present in the broader environment, as well as general protection against the diverse array of antimicrobial metabolites known to be produced by lichens (Ranković *et al*. 2007; Studzińska-Sroka *et al*. 2015; Taylor *et al*. 2023). While there is currently no evidence that *C. stellaris* specifically synthesizes quinolone-type compounds, the selective pressure exerted by its secondary metabolite profile could favor the retention of versatile detoxification mechanisms like *qepA* in the lichen-associated bacteria. This evolutionary retention would explain both the prevalence and exceptional abundance of *qepA* in *C. stellaris*. Network analysis revealed that *Tundrisphaera* and *Terriglobus* were positively associated with *qepA*, although both genera exhibited low betweenness centrality within the bacterial network. This suggests that despite being carried on MGEs (Jacoby *et al*. 2006; Périchon *et al*. 2007), *qepA* may not disseminate widely across the lichen microbiome due to the limited ecological centrality of its primary bacterial hosts.

#### Occasional ARGs: aminoglycosides, sulfonamides, and int1-a-marko

In addition to the dominant ARGs discussed above, we detected the occasional presence of other resistance genes and the integron marker gene *int1-a-marko* in *C. stellaris*. Specifically, the aminoglycoside resistance genes *aac(6’)-Ib* and *aac(3’)*, the sulfonamide resistance gene *sul1*, and the integron marker gene *int1-a-marko* were identified at low frequencies, predominantly in samples from the southern LW. Aminoglycosides are natural compounds produced by soil bacteria (Schatz *et al*. 1944), and resistance to these compounds is widespread across diverse natural environments (Li *et al*. 2015; Yan *et al*. 2017; Zhuang *et al*. 2021; Urban-Chmiel *et al*. 2022). In contrast, sulfonamides are synthetic antibiotics first introduced in the 1930s, widely used in both human and veterinary medicine (Landers *et al*. 2012), with resistance genes now also broadly disseminated in the environment (Nunes *et al*. 2020). The geographic clustering of these ARGs in southern LW samples suggests that anthropogenic influence from nearby urban areas, potentially mediated by bioaerosol dispersal, could have introduced them into the PNGJ. However, the very limited number of positive detections makes it difficult to support this hypothesis alone, and alternative explanations, such as co-evolution, cannot be excluded. Additionally, we detected *sul2* in more than 40% of samples from both northern and southern regions. The frequent detection of *sul2* may indicate widespread resistance. Resistance to sulfonamides has been documented in bioaerosols (Zhang *et al*. 2018; Han and Yoo 2020; Qu *et al*. 2024). However, it has also been detected in forest soils with no history of exposure to these synthetic drugs (Willms *et al*. 2019). This suggests that, although sulfonamides may not naturally occur in these environments, the genetic mechanisms for resistance can still persist within microbial communities. Such persistence indicates an adaptive capacity that may explain *sul2* persistence within the *C. stellaris*, despite limited exposure to antibiotics from anthropogenic uses.

## Conclusion

To our knowledge, this is the first study to directly quantify and confirm the presence of ARGs in lichens using high-throughput quantitative PCR. We detected ten ARGs and one MGE in the lichen *C. stellaris* from northern and southern LWs in eastern Canada, with predominance of beta-lactam and quinolone resistance genes. Our results revealed differences in ARG relative abundances between northern and southern sites, although only *qnrB* showed significant variation. Furthermore, the bacterial community composition, while distinct between LWs, did not appear to drive ARG distribution. Two hypotheses were proposed to explain the presence and distribution of the most prevalent ARGs detected in *C. stellaris* (*blaCTX-M-1*, *qnrB*, and *qepA*): long-distance dispersal via bioaerosols and, endogenous development through co-evolutionary dynamics between lichens and their associated bacteria. We hypothesize that *blaCTX-M-1* and *qepA* may reflect ancient co-evolutionary processes, while the higher abundance of *qnrB* in southern samples may suggest a more recent introduction linked to anthropogenic influence. However, given the exploratory nature of this study and the complexity of ARG dynamics in natural systems, these interpretations remain to be tested. Moreover, the specific atmospheric processes potentially contributing to ARG dispersal, such as wildfire smoke, prevailing winds, precipitation, or cloud dynamics, were not assessed and remain to be investigated in future research. As antibiotic resistance continues to pose a major global health challenge, understanding how resistance genes persist and spread even in isolated environments such as LWs represents an open and relevant scientific question.

## Author contributions

**Marta Alonso-García**: Conceptualization; Funding acquisition; Investigation; Methodology; Data analysis; Writing – original draft; Writing – review & editing. **Paul George**: Funding acquisition (supporting); Data analysis (supporting); Writing – review & editing. **Samantha Leclerc**: Investigation (supporting); Methodology (supporting). **Marc Veillette**: Supervision (laboratory coordination); Methodology (advice on experimental procedures). **Caroline Duchaine**: Funding acquisition (supporting), Supervision; Writing – review & editing. **Juan Carlos Villarreal**: Conceptualization; Funding acquisition, Supervision; Writing – review & editing.

## Conflict of interest

The authors declare that the research was conducted in the absence of any commercial or financial relationships that could be construed as a potential conflict of interest.

## Data availability

The 16S rRNA sequencing data analyzed in this study were generated in a previous project and are publicly available in the NCBI Sequence Read Archive under BioProject accession numbers PRJNA593044. Detailed sample metadata, including BioSample accession numbers, geographic coordinates, and collection details, are provided in Table S1.

## Acknowledgements

This research was supported by the Sentinel North Research Acceleration Fund (Laval University), the CRNSG-RGPIN/05967-2016, and the *Fondation canadienne pour l’innovation*. The original sample collection was made possible thanks to the *Centre d’étude nordique*, which provided research infrastructure and logistical support in Kuujjuarapik-Whapmagoostui, and to the authorities of *Parc national des Grands-Jardins* (SEPAQ) for granting permission. We are grateful to the staff of *Herbier Louis-Marie* for their collaboration. We also thank the *Plateforme d’analyse génomique* and the *Plateforme de bio-informatique* of the *Institut de biologie intégrative et des systèmes* (Université Laval) for their assistance during the initial stages of data acquisition and analysis.

## Supplementary data

Supplementary data are available at Annals of Botany online and consist of the following:

**Table S 1**. Detailed information on the *Cladonia stellaris* samples analyzed in this study and collected from northern lichen woodlands in Kuujjuarapik-Whapmagoostui and southern lichen woodlands in Parc National des Grands-Jardins (PNGJ). Each entry includes the voucher number, collector’s name, collection number, geographic position, specific collection sites, altitude range, and precise latitude and longitude coordinates. Additionally, for each sample, the BioSample accession number (Accession) and associated BioProject code (BioProject) registered in the NCBI database are provided.

**Table S 2**. List of bacterial genera identified in *Cladonia stellaris* samples with their respective ASV counts.

**Table S 3**. Relative abundance of antibiotic resistance genes (ARGs) and a mobile genetic element (MGE) in *Cladonia stellaris* from northern and southern lichen woodlands (LWs). The columns list the specific ARGs detected, and the rows represent individual lichen samples. The LW from which each sample originates is indicated. The numeric values indicate the abundance of each ARG/MGE relative to the 16S rRNA gene, with values greater than 1 signifying a higher abundance than 16S rRNA, and values less than 1 indicating a lower abundance. “NA” denotes that the gene was not detected in the corresponding sample.

**Table S 4**. Summary of network metrics for bacterial genera and antibiotic resistance genes (ARGs) in *Cladonia stellaris* from northern and southern lichen woodlands (LWs). The table presents detailed network statistics, including degree, closeness centrality, betweenness centrality, clustering coefficient, average shortest path length, eccentricity, neighborhood connectivity, number of directed and undirected edges, stress, radiality and topological coefficient. Metrics are shown for all samples. For bacterial genera, additional columns include the phylum, class, order, family and genus information, while ARGs are listed with their specific resistance mechanisms. When the genus is unknown, the bacteria are identified by the family name followed by’Unclassified’.

**Figure S 1.** Relative abundance per phylum (**A**) and genera (**B**) of bacteria associated to *Cladonia stellaris*. Samples are grouped by latitude of lichen woodland (LW), northern or southern, as indicated at the top of the bar plot. Each vertical bar represents a single sample, and colors reflect different phyla (**A**) or genera (**B**). In panel B, unassigned taxa were removed prior to plotting, which may result in bar heights summing to less than one.

**Figure S 2.** Total relative abundance of ten antibiotic resistance genes (ARGs) and a mobile genetic element (MGE) in *Cladonia stellaris* from northern and southern lichen woodlands (LWs). The boxplots depict the median (line in the box), quartiles (edges of the box), and range of values within 1.5 times the interquartile range (whiskers) of the 11 ARGs/MGE within each LW. Individual points represent individual observations. The y-axis is on a logarithmic scale to better visualize differences. Wilcoxon rank-sum results indicate no significant differences between the northern and southern LWs (adjusted p-value = 0.3669).

## Notes

### Competing Interest Statement

The authors have declared no competing interest.

