## Supplementary figures and images for "Antibiotic resistance genes detected in lichens: insights from *Cladonia stellaris*"

### Figure S1A

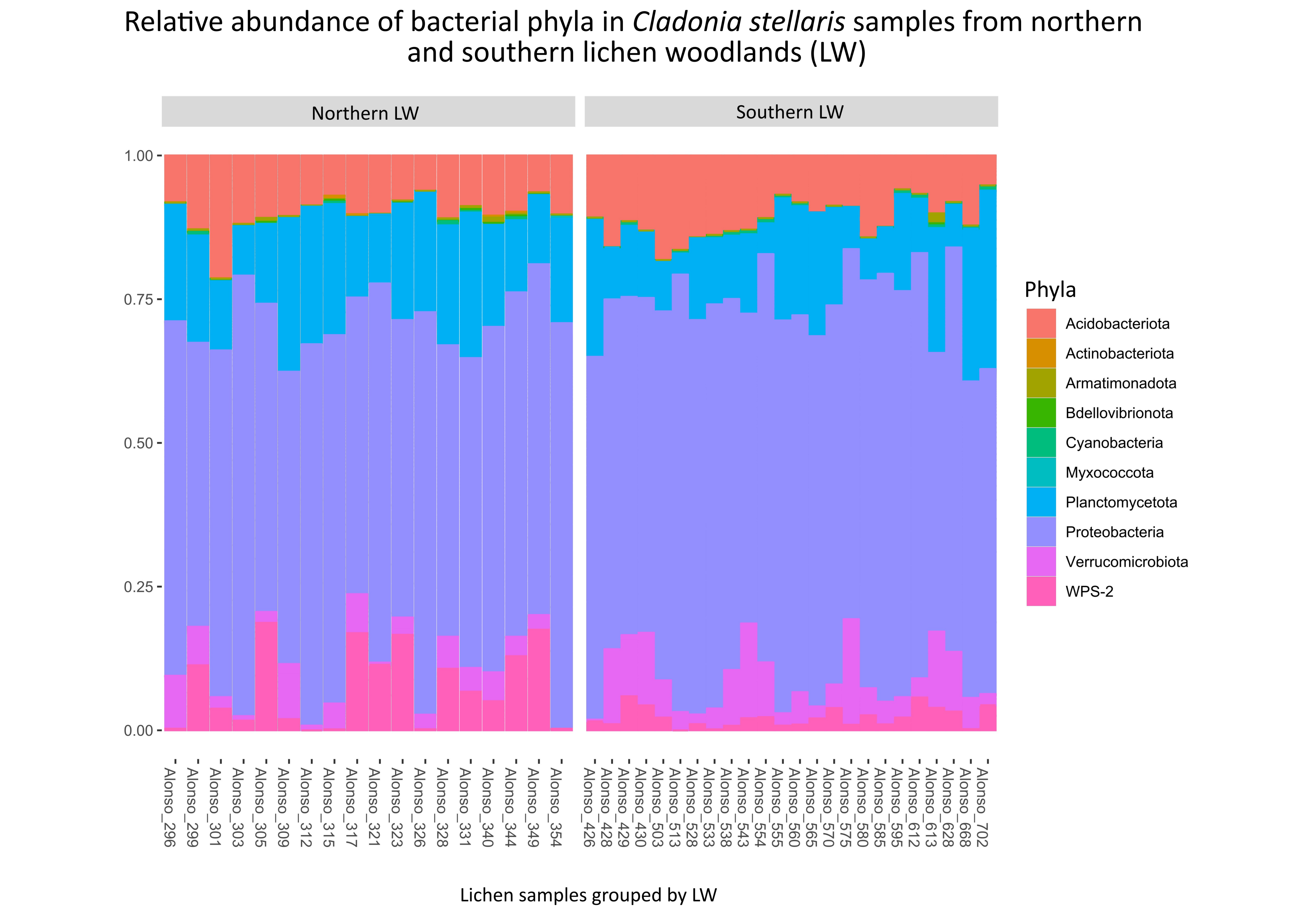

### Figure S1B

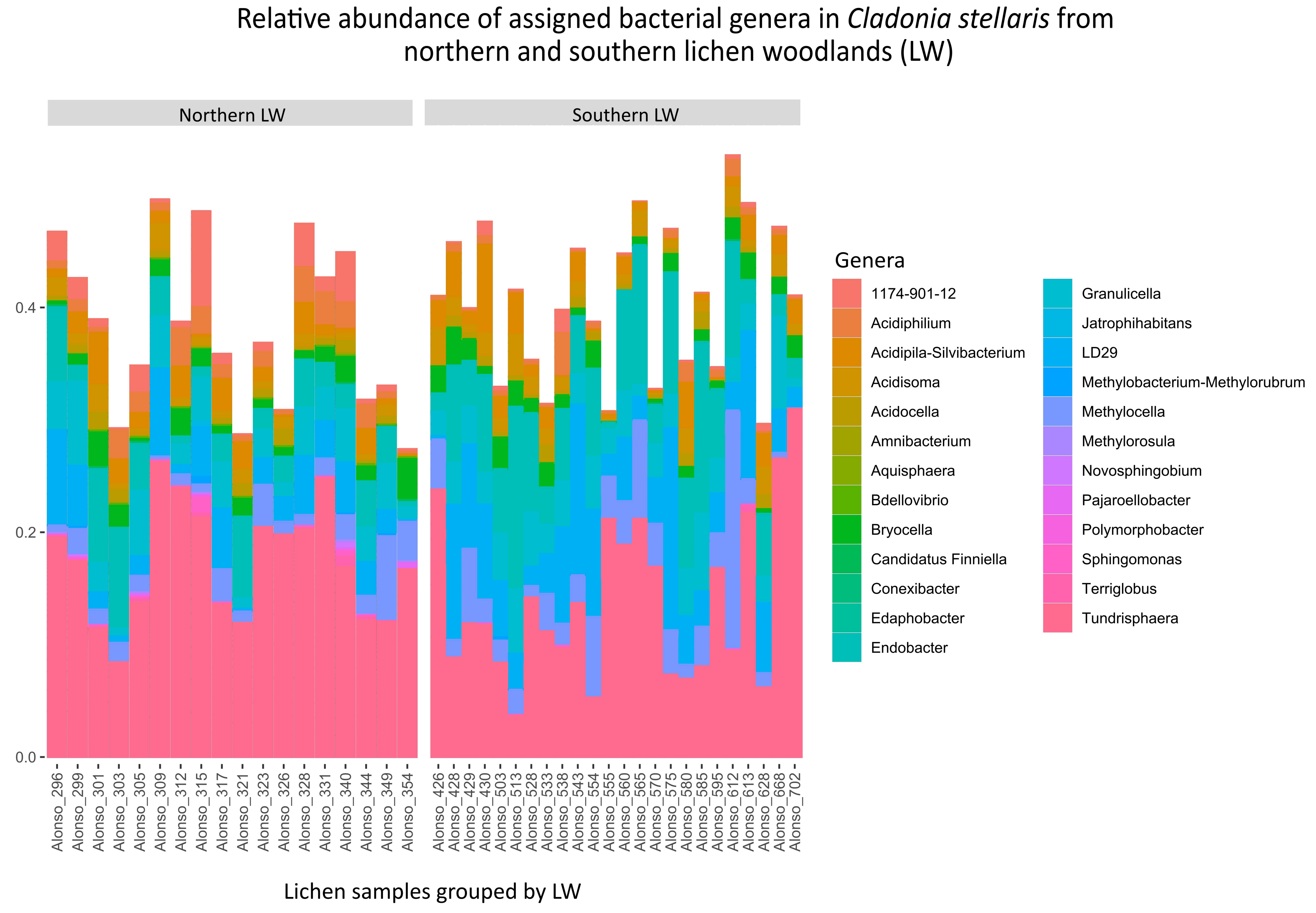
