## Supplementary material for "Antibiotic resistance genes detected in lichens: insights from *Cladonia stellaris*": Figure S2

### Comparison of total relative abundance of antibiotic resistance genes (ARGs) and mobile genetic element (MGE) from northern and southern lichen woodlands (LW)

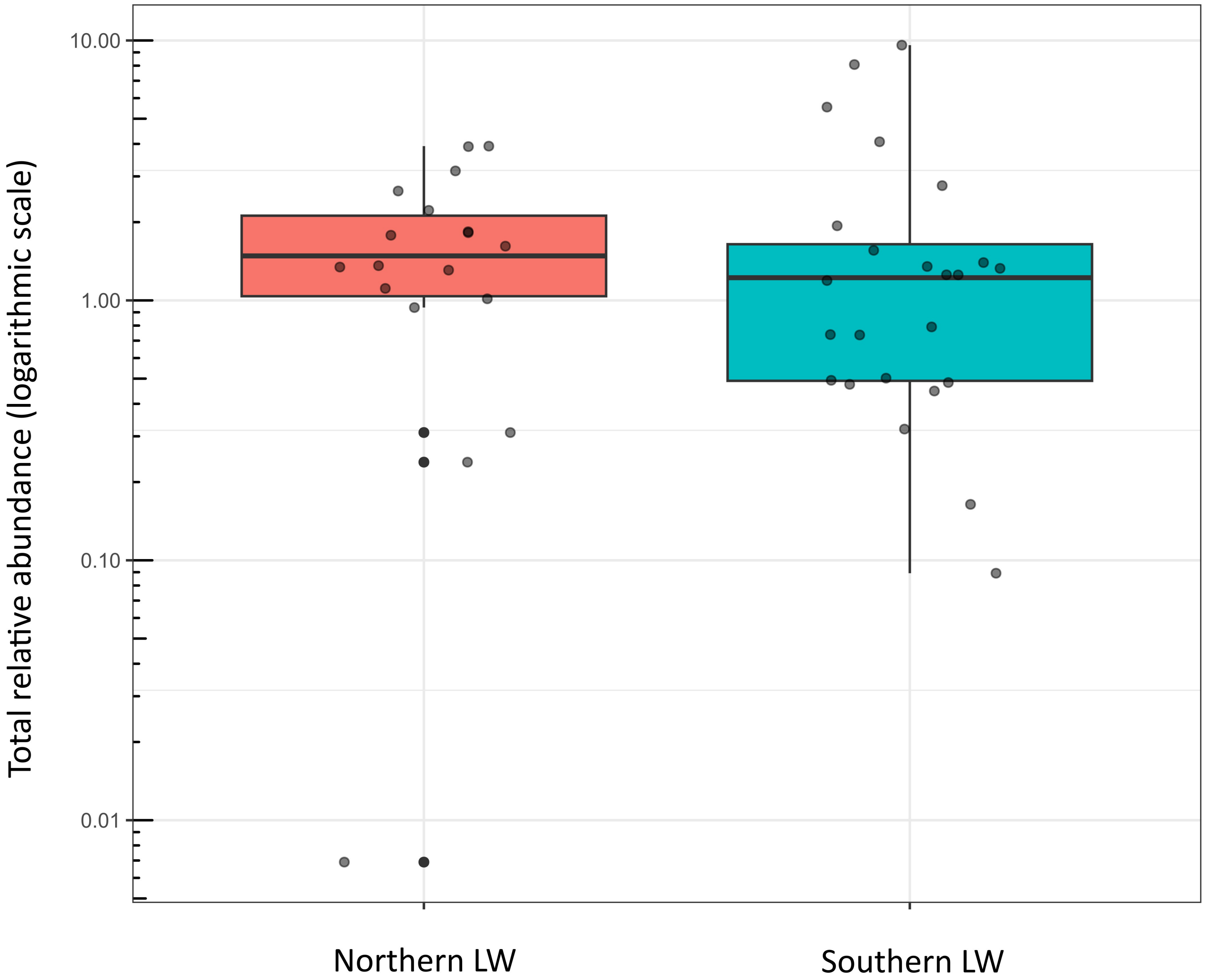
